# Did *Moldehawera* flowers evolve through mimicry with oil-producing Malpighiaceae?

**DOI:** 10.1101/2023.06.15.545073

**Authors:** Luciano Paganucci de Queiroz, Jorge Antonio Silva Costa, Cristiana Barros Nascimento Costa

## Abstract

Floral mimicry is a captivating phenomenon wherein flowers imitate traits of other species to attract specific pollinators. The Caesalpinioideae legumes in general have relatively unspecialized flowers, which has allowed the development of disparate morphologies associated with adaptation to different types of pollinators. This study describes the pollination of *Moldenhawera nutans* and explores its potential floral mimicry towards Malpighiaceae flowers. Our investigation revealed that *M. nutans* is pollinated by species of *Centris* Fabricius, 1804 and *Xylocopa* Latreille, 1802. It also presents compelling evidence supporting the hypothesis of floral mimicry, including striking similarities in floral display, shared oil-collecting pollinators, oil collection behavior in *M. nutans* despite the absence of oil production, and the reliance on exogenous pollen for reproduction. These findings suggest that species of *Centris* bees visit *M. nutans* flowers under the mistaken impression of oil availability, subsequently transitioning to pollen collection. We explored other potential cases of floral mimicry with Malpighiaceae in the Caesalpinioideae legumes by optimizing the Malpighiaceae-like floral display on a dated phylogeny of this subfamily. However, current information does not allow us to determine whether the similarities in floral morphology represent cases of floral mimicry, phylogenetic inertia, or simple convergence. Several hypothesis tests are suggested that can guide the study of these fascinating evolutionary processes in the group.

## Introduction

Among the strategies employed by flowering plants to enhance their reproductive success through biotic pollination is floral mimicry, a fascinating adaptive strategy in which flowers develop traits that resemble those of other species, thereby attracting specific pollinators. It involves the convergent evolution of “common advertising displays” in flowers of unrelated species, enabling the mimic species to exploit the specialized pollinators of the mimetic models (Macior 1971; Proctor and Yeo 1972; Brown and Kodric-Brown 1979; Thomson 1981; Schemske 1981; Little 1983; Dafni 1984; Roy and Widmer 1999).

*Moldenhawera* Schrad. is a small genus of the subfamily Caesalpinioideae (Leguminosae) confined to eastern and northeastern Brazil. It comprises twelve species of trees or shrubs primarily inhabiting humid forests, restingas (shrublands and low coastal forests on sandy soils), and rupestrian fields on sandy soils in the Espinhaço Range mountains (Queiroz et al. 1999; Vivas et al. 2015, 2019; Vivas and Queiroz 2020). The flowers are pentamerous (rarely tetramerous) and possess a distinct floral morphology that is conserved across the genus (Figure 1). The flowers exhibit slight bilateral symmetry. The sepals are separate and reflexed when the flower opens, revealing the bright green or yellow color on the inner surface of the sepals. The bright yellow petals have a broad and wrinkled blade with wavy or fringed margins and a long claw, except for *M. acuminata* Afr. Fern. & P. Bezerra, which has light pink or white petals. The androecium consists of eight stamens (in tetramerous flowers) or ten stamens (in pentamerous), but only one stamen is fertile. This fertile stamen bears an anther with longitudinal dehiscence and a long filament, and, together with the style, it is displaced towards one side of the flower, aligning the anther at the same level and close to the stigma. The remaining stamens are staminodes and possess very short filaments, with their anthers barely exceeding the length of the ovary. The anthers of staminodes typically dehisce through two apical pores, although in *M. acuminata*, they open via two small median slits, and in *M. papillanthera* L.P. Queiroz, G.P. Lewis & Allkin they do not open at all.

**Figure 1.**
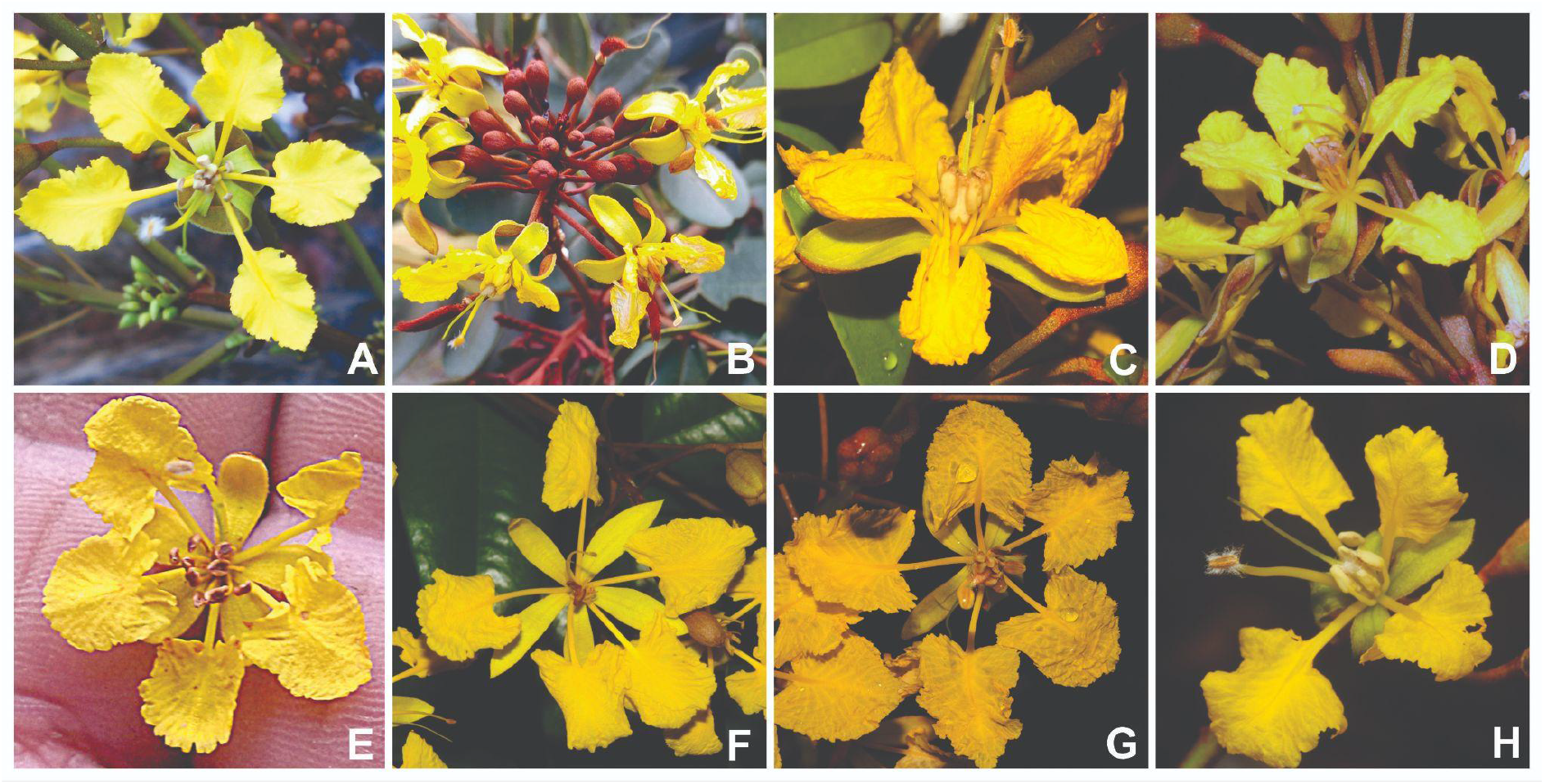
Flowers of different species of *Moldenhawera*. **A**, *M. blanchetiana* Tul.; **B**, *M. brasiliensis* Yakovlev; **C**, *M. congestiflora* C.V. Vivas & L.P. Queiroz; **D**, *M. emarginata* (Spreng.) L.P. Queiroz & Allkin; **E**, *M. floribunda* Schrad.; **F**, *M. longipedicellata* C.V. Vivas & L.P. Queiroz; **G**, *M. luschnathiana* Yakovlev; **H**, *M. nutans* L.P. Queiroz, G.P. Lewis & Allkin. Photos. A: Lucas Marinho; B: Rubens Queiroz; C, D, G, Luciano P. Queiroz; E: Alex Popovkin; F: Cláudio N. Fraga; H: Domingos Cardoso.

The flowers of *Moldenhawera* species bear a striking resemblance to those of neotropical species in the Malpighiaceae family. This family comprises approximately 1250 species, with nearly 1000 species found in the Americas and the rest distributed in Africa and a few Asian species (Anderson 1990). The American species inhabit a wide range of habitats and exhibit significant morphological diversity in terms of their growth habit, leaves, and fruits, while maintaining a remarkably similar floral structure (Anderson 1979, 1990). The flowers display slight zygomorphy, with the five or four sepals typically possessing two prominent elaiophores (oil-secreting glands) on their outer surfaces (Vogel 1974, 1990; Anderson 1979). The five separate petals have a wide blade and a conspicuous claw. Among them, the flag petal usually differs in size and possesses a more robust claw. The petals often reflex between the sepals, facilitating easy access to the calyx glands for insects that land in the center of the flower.

The main pollinators of neotropical Malpighiaceae are specialized oil-collecting Centridini bees (Vogel 1974, 1990; Anderson 1979, 1990) while the African and Asian species do not produce oils, representing at least six losses of the floral oil secreting glands in Old World clades (Davis et al. 2001, 2004). The oil-collecting behavior involves the positioning of the bee in the center of the flower that then grasps the claw of the flag petal with its mandibles, and then scrapes the calyx glands using modified blade-like setae of its fore and midlegs (Sazima and Sazima 1989). The collected fatty oil mixed with pollen are used as food for the larvae. The presence of oil-glands on the sepals is restricted to the American species of Malpighiaceae, although they are not universal. Some American species are eglandular and there may be intraspecific variation in the presence of elaiophores (Sazima and Sazima 1989). Anderson (1979) considered that the floral syndrome adapted to specialized reward to a restricted group of oil collecting bees as pollinators in neotropical Malpighiaceae “explains why the flowers have remained so conservative in spite of the evolution of great diversity in other aspects of the phenotype” (Anderson 1979: 219).

This study focuses on investigating the hypothesis of floral mimicry in *Moldenhawera nutans* L.P. Queiroz, G.P. Lews & Allkin in relation to Malpighiaceae flowers. Supporting this hypothesis, we observe striking resemblances in floral display between *M. nutans* and Malpighiaceae species (Figure 2) and the sympatric occurrence of both of them along their distribution areas could indicate that these similarities involve mechanisms of mimicry.

**Figure 2.**
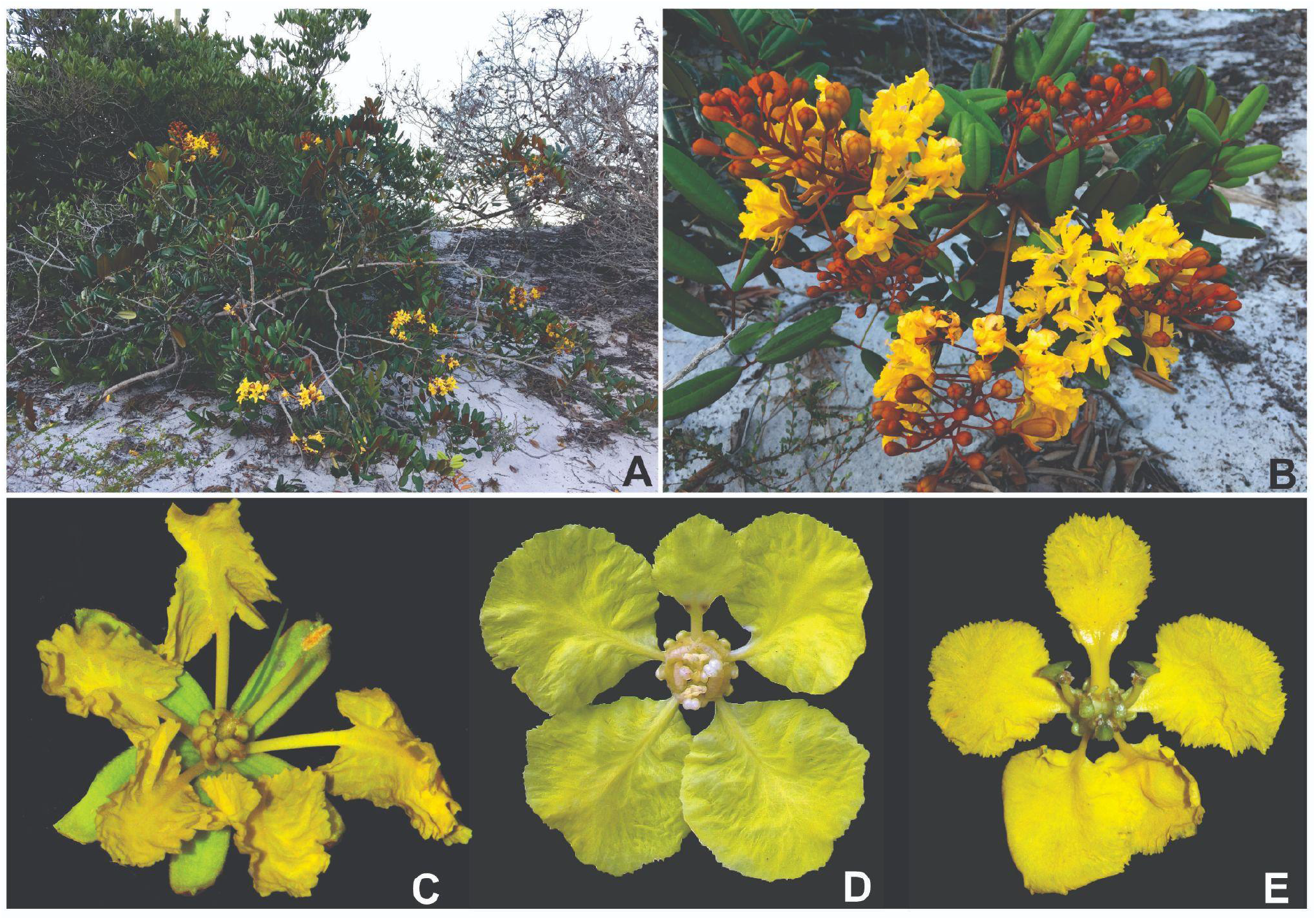
*Moldenhawera nutans* L.P. Queiroz, G.P. Lewis & Allkin in its natural habitat in coastal sand dunes of Salvador municipality, Bahia state (**A**) and a detail of a flowering branch (**B**). The Malpighiaceae-like flower of *M. nutans* (**C**) and flowers of *Peixotoa hispidula* A. Juss. (**D**) and *Stigmaphyllon paralias* A. Juss. (**E**), two sympatric species of Malpighiaceae that were found blooming at the same time as *M. nutans*. Sources of pictures: A-C: Domingos Cardoso; D: Rafael F. Almeida; E: Marco Pellegrini.

Additionally, we investigated the evolution of Malpighiaceae-like flowers within the grade of the Caesalpinioideae subfamily that subtends the mimosoid clade. Flowers in this grade are relatively unspecialized with radial or weakly bilateral symmetry, free sepals, petals, and stamens, and limited nectar protection, allowing for the evolution of a wide range of floral morphologies in response to different groups of pollinators (Arroyo 1981), even within more restricted lineages (Gagnon et al. 2016). Malpighiaceae-like flowers occur in different genera of the Caesalpinioideae subfamily, as reported by Sazima et al. (2009) for *Schizolobium parahyba* (Vell.) S.F. Blake. These genera appear scattered in the recently published phylogeny of the subfamily by Ringelberg et al. (2022), suggesting multiple cases of convergence within the Caesalpinioideae and we investigated whether they could be considered additional cases of potential floral mimicry with Malpighiaceae.

## Material and methods

Observations on the floral biology of *Moldenhawera nutans* were carried out in two neighboring areas in the municipality of Salvador. JASC and CBNC carried out observations in the Parque Metropolitano do Abaeté region, in the Stella Maris neighborhood (hereinafter Site 1; 12°56’26”S, 38°20’56”W) between October 29 and November 14, 2001. LPQ carried out the same studies at the dunes around the Urubu lagoon, in the Alamedas da Praia neighborhood (hereinafter Site 2; 12°55’24”S, 38°20’40”W) between January 18 and 22, 2016. Both areas include restinga vegetation over coastal sand dunes. The physiognomy of the vegetation is predominantly shrubby in thickets separated by open areas of bare sand (Figure 2).

*Moldenhawera nutans* is a bushy plant with 1.5 to 2.5 m high, quite branched, with spreading and sometimes almost horizontal branches, forming rounded clumps. The flowers are grouped in erect terminal panicles that leave the flowers exposed above the foliage. The flowering of *M. nutans* it is concentrated between October and February (Queiroz et al. 1999; Viana et al. 2006).

Two species of Malpighiaceae with flowers similar in size and color to those of *M. nutans* were observed to occur sympatrically and blooming at the same time as this species. *Stigmaphyllon paralias* A. Juss. is an erect subshrub 0.3 to 1 m tall, usually occurring in dense populations around the thickets formed by *M. nutans* at Sites 1 and 2. The second species is *Peixotoa hispidula* A. Juss., a vine that climbs in bushes in the thickets, observed only in Site 2. The flowering of these two species occurs throughout the year (flowering data from *S. paralias* were presented by Costa et al. 2006; Viana et al. 2006, while those of *P. hispidula* were extracted from specimen images available online on the website of the Species Link).

Field observations started around 06:00 am and were concentrated in the morning. Floral visitors were observed regarding foraging behavior and floral resources collection. Specimens of bee species visiting flowers of the focal plant species were collected and after identified were deposited at the Botany Laboratory of the Sosígenes Costa Campus of the Federal University of Southern Bahia. Flower buds were bagged with voile bags to verify the occurrence of agamospermy and spontaneous self-pollination.

In addition to reporting a potential example of floral mimicry in *Moldenhawera*, we aimed to investigate the evolution of the Malpighiaceae-like floral display in the Caesalpinioideae subfamily. Malpighiaceae-like flowers were defined by the following syndrome of morphological characters: radial or weakly bilateral symmetry, yellow clawed petals with a wide and wrinkled lamina, and wavy or indented margins forming an open corolla, without protection for floral resources, thus making them accessible to different types of bee pollinators. Therefore, not all flowers of Caesalpinioideae species with yellow clawed petals were considered as Malpighiaceae-like. Several species within the Caesalpinia clade (sensu Ringelberg et al. 2022) were not considered as having this floral display because their symmetry is clearly bilateral, and access to nectar is protected by the position of the adaxial petal. In the Cassia clade (sensu Ringelberg et al. 2022), species of *Cassia, Senna*, and *Chamaecrista*, also with yellow petals, were coded as not presenting a Malpighiaceae-like floral display because they have flowers adapted for buzz-pollination by large carpenter bees (Buchmann 1983; Westerkamp 2004).

To investigate the evolution of the Malpighiaceae-like floral display in the Caesalpinioideae subfamily, we encoded it as a binary character (present/absent) and optimized its presence on a recently published dated phylogeny of the Caesalpinioideae subfamily by Ringelberg et al. (2023), based on 997 nuclear genes. Its evolutionary history was then evaluated using the “Trace character history” option in Mesquite v. 3.81, using the Markov k-state 1-parameter likelihood model (Maddison & Maddison 2023). We then compared the obtained dates for the origin of lineages with the Malpighiaceae-like floral display with the estimated ages for the origin of oil-producing Malpighiaceae and oil-collecting Centridini bees. For Malpighiaceae, we used the estimated age range between the average ages of the crown group (59.8 Ma, 95% HPD 69.0–52.5 Ma), and the stem group (86.1 Ma, 95% HPD 99.9–72.9 Ma; Xi et al. 2012). For the Centridini bees, Martins et al. (2014) estimated the origin of the oil-collecting behavior in the Centridini bees (the Apini clade) at 91 Ma (95% HPD 103–79 Ma). However, since the Centridini form a paraphyletic group with respect to corbiculate bees, it is possible that the oil-collecting behavior evolved independently in the two Centridini lineages, the genera *Epicharis* Klug, 1807 and *Centris*. In the case of *Centris*, which includes species reported to visit *Moldenhawera* flowers, the origin of the genus (crown age) was estimated at 46 Ma (95% HPD 59-36 Ma; Martins et al. 2014).

## Results

### Floral morphology of *Moldenhawera nutans, Stigmaphyllon paralias* and *Peixotoa hispidula*

The flowers of *M. nutans* have a diameter ranging from 38.7 to 39.0 mm. When open, the sepals reflex, revealing their bright green color. The petals are a golden yellow color, clawed, with a wide ovate to suborbicular wrinkled blade and a margin that is indented to fringed. The fertile stamen measures 19 to 23 mm in length, with a densely wooly connective, and the anther opens through two longitudinal slits. The staminodes are positioned centrally in relation to the open flower’s axis, surrounding the ovary, and they measure 4 to 5 mm in length, with anthers that open through two apical pores. The style is similar in length to the filament of the fertile stamen, resulting in the stigma and anther being very close together and at the same height, slightly displaced to one side of the flower (Fig. 2C).

None of the bagged flowers of *M. nutans* resulted in fruit formation, demonstrating that in this species, neither self-pollination nor apomixis occurs, emphasizing the requirement for exogenous pollen for reproduction.

The flowers of *Stigmaphyllon paralias* have an average diameter of 24.09 ± 1.73 mm (n = 10 flowers). Most flowers possess two elaiophores on each sepal, totaling 10 elaiophores per flower, while some flowers may have one sepal lacking elaiophores, resulting in a total of 8 elaiophores. The petals are golden yellow and clawed, with the flag petal displaying a slightly larger size and a thicker claw compared to the other petals. The ten fertile stamens are relatively short (approximately 5 mm in length) and cluster around the ovary in the center of the flowers. The ovary is centrally positioned within the flower and is topped by three wide styles (Fig. 2D; Almeida 2020b).

The flowers of *Peixotoa hispidula* closely resemble those of *S. paralias*. They have a diameter of 32.17 ± 2.05 mm (n = 10 flowers). All observed flowers possess 8 elaiophores, with one sepal lacking elaiophores. The flag petal is slightly smaller and has a thicker claw. The androecium consists of five fertile stamens and five staminodes, clustered around the ovary at the center of the flower (Fig. 2E; Almeida 2020a). Due to the presence of the flag petal, both species of Malpighiaceae exhibit slight zygomorphy in their flowers.

Despite some differences in detail, the flowers of *M. nutans, S. paralias*, and *P. hispidula* exhibit striking similarities in their overall architecture and appearance. They all display slightly zygomorphic flowers with yellow clawed petals featuring a suborbicular wrinkled and fringed blade. The male reproductive structures are clustered in the center of the flower around the ovary, with *M. nutans* having staminodes and the two Malpighiaceae species having stamens or a combination of stamens and staminodes. Furthermore, the cylindrical anthers of the staminodes, clustered around the ovary, resemble the elaiophores of Malpighiaceae flowers when the flower of *M. nutans* is observed from the front, further enhancing the similarity in the floral display.

### Flower visitors and visiting behavior

Bees of the genera *Centris* (Apidae Centridini) and *Xylocopa* (Apidae Xylocopini) have been observed making legitimate visits to the flowers of *M. nutans*, as previously recorded by Viana and Kleinert (2006). At Site 2, *Trigona spinipes* (Fabricius, 1793) (Apidae Meliponini) was observed making illegitimate visits to the flowers of *M. nutans*, typically cutting the staminode anthers and collecting pollen without contacting the anther or stigma (Table 1). Three species of *Centris*, namely *Centris sponsa* Smith, 1854, *C. flavifrons* Fabricius, 1775, and *C. leprieuri* Spinola, 1841, were observed visiting the flowers of *M. nutans* and Malpighiaceae species. *Centris sponsa* was observed at both sites, while *C. flavifrons* and *C. leprieuri* were only observed at Site 1.

**Table 1.**
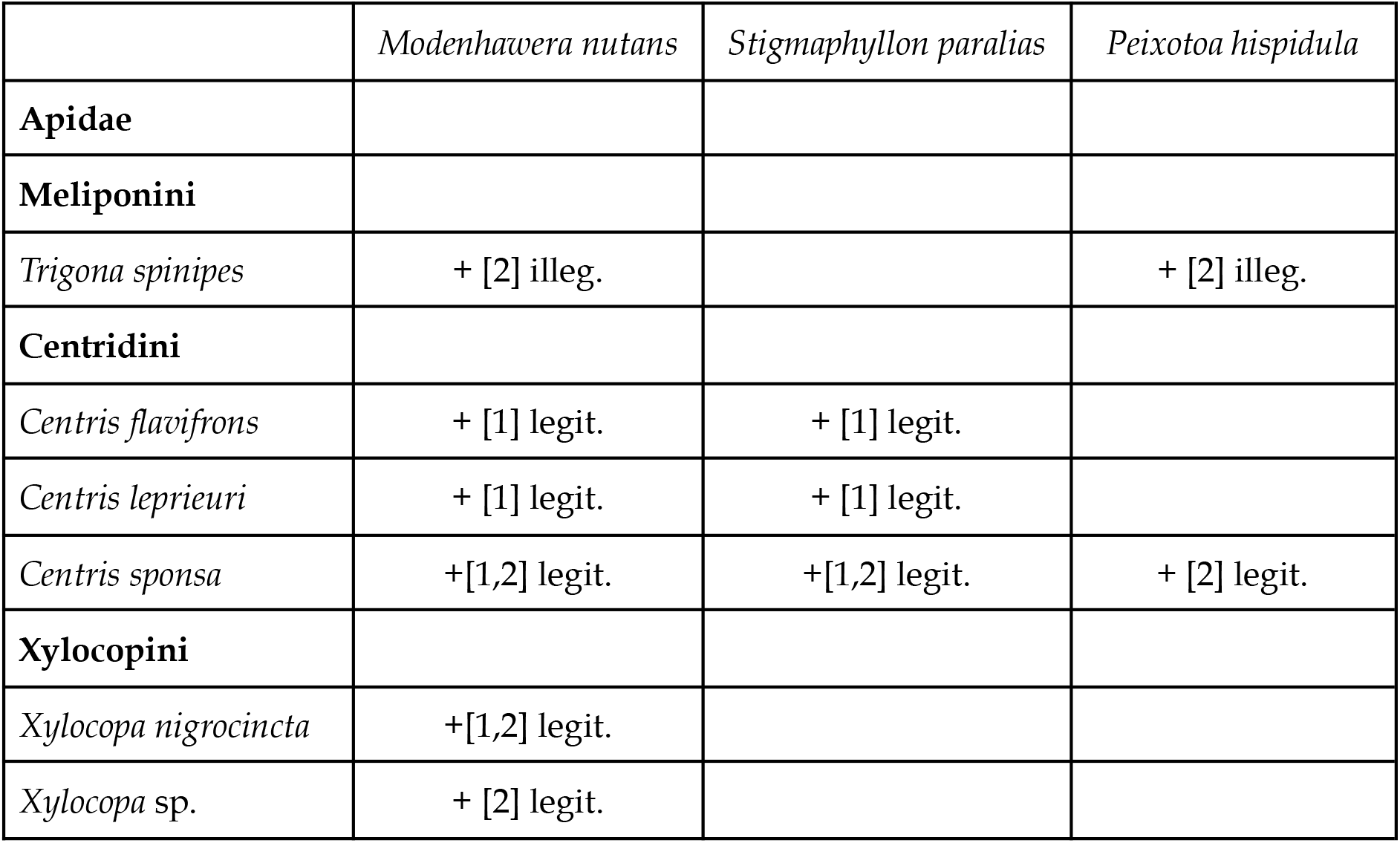
Floral visitors observed in *Moldenhawera nutans* (Leguiminosae), *Peixotoa hispidula* and *Stygmaphyllon paralias* (Malpighiaceae). The “+” sign indicates that the visit of the bee species was observed. The number in square brackets indicates the Site where the interaction was observed. Legitimate visits are indicated by “legit.” and illegitimate ones by “illeg.”

*Centris* species exhibit two types of visits to the two species of Malpighiaceae. The most common visits involve collecting the oil produced in the elaiophores. During these visits, the bees land at the center of the flower, facing the flag petal. They then use their mandibles to grasp the claw of the petal and apply upward pressure on the elaiophores using their fore and mid legs. After approximately 10 seconds, the bee moves on to another flower. This oil collection behavior is identical to that observed in other Centridini bee species visiting various Malpighiaceae species in southeastern Brazil (Sazima and Sazima 1989; Sigrist 2001; Sigrist and Sazima 2004; Sazima et al. 2009).

*Centris* species also engage in another type of visit to Malpighiaceae species, which involves gathering pollen. During this visit, the bees cluster their fore and mid legs around the calyx, extend their hind legs, and vibrate the flower. The behavior of pollen collection in Malpighiaceae species was described by Sazima and Sazima (1983), where different *Centris* species visit both glandular and eglandular morphs of *Banisteriopsis muricata* (Cav.) Cuatrec. for the collection of oil or pollen.

The same species of *Centris* were observed visiting flowers of *M. nutans*. At Site 1, visits by *C. sponsa* occurred between 06:00h and 10:00h, followed by *C. flavifrons* and *C. leprieuri* after 10:00h until 12:00h. At Site 2, the only species of *Centris* observed visiting *M. nutans* was *C. sponsa*, until approximately 09:30h. The visiting behavior of these *Centris* species in *M. nutans* initially resembled the oil collection behavior in Malpighiaceae flowers. After landing in the center of the flower, the bees “embraced” the flower and performed upward movements with their legs over the sepals, as if they were attempting to extract oil. Upon realizing the mistake, the bees quickly (<10 seconds) changed their behavior, extended their hind legs, and collected pollen through vibration. While collecting pollen from the staminodes, the bees made contact with the anther and stigma in the dorsolateral region of the thorax, indicating their effectiveness as pollinators. This change in behavior from oil collection to pollen collection within the same flower was also reported by Sazima and Sazima (1989) in eglandular morphs of *Banisteropsis muricata*. The use of mandibles to grasp the flag petal was not observed, a behavior also noted by Costa et al. (2006) in a study on the reproductive biology of five Malpighiaceae species conducted in Site 1.

*Xylocopa nigrocincta* Smith, 1854 was observed at Site 1, and an unidentified species of *Xylocopa* was observed at Site 2, visiting the flowers of *M. nutans*. The *Xylocopa* species only collected pollen through vibration and did not exhibit oil collection behavior. Due to their position within the flower of *M. nutans* and their large size, they made lateral contact with the anther and stigma on the thorax, indicating their effectiveness as pollinators of this species. The visitation of these two *Xylocopa* species to the flowers of the Malpighiaceae species was not observed.

The presence of *Centris* species as effective pollinators of both *M. nutans* and sympatric Malpighiaceae species, along with their behavior in the flowers, is comparable to what was reported by Sazima and Sazima (1989) in glandular and eglandular morphs of Malpighiaceae species. In the case of glandular morphs, *Centris* species visit the flowers and collect oil using the described manner. However, when visiting eglandular morphs, the bees initially attempt to collect oil but quickly switch to pollen collection through vibration. This similarity is evident in both *M. nutans* and the other Malpighiaceae species, suggesting that the bees are unable to distinguish between glandular and eglandular flowers. As a result, oil-collecting bees can still contribute to the pollination of eglandular flowers.

### Do the flowers of *Moldenhawera nutans* mimic those of Malpighiaceae?

The hypothesis that the flowers of *M. nutans* mimic the flowers of Malpighiaceae is supported by several pieces of evidence. There is a strong similarity in the floral display between them in shape and color, as well as the fact that they share the same species of oil-collecting pollinator bees. The importance of floral display similarity has been indicated in several studies, which demonstrate that this similarity can occur in different ways, including shapes, colors, and/or odors, depending on the biology of shared pollinators (Johnson 1994; Johnson et al. 2003; Peter et al. 2008; Papadopulos et al. 2013). Furthermore, the oil-collecting behavior in the flowers of *M. nutans*, despite not producing this reward, and the reliance on external pollen for reproduction provide additional support. These observations suggest that species of *Centris* visit the flowers of *M. nutans* mistakenly and only switch to pollen collection once they realize their error, a behavior also observed in eglandular flowers of typically glandular species of Malpighiaceae (Sazima and Sazima 1989). These are four out of the five criteria listed by Roy and Widmer (1999) for the similarity between two or more species to be considered a case of floral mimicry. The fifth criterion, which states that the similarity should be important for fitness, was not tested in this study (see discussion below regarding potential tests for this criterion).

Two types of floral mimicry are recognized. In Batesian floral mimicry, a non-rewarding species mimics a model species that provides some form of reward to the pollinator, such as nectar or pollen. In Müllerian floral mimicry, both species involved provide floral rewards to the same pollinator species, resulting in strong convergence towards a similar floral display. In the case of *M. nutans* and Malpighiaceae species, there appears to be an intermediate situation between these two types of mimicry. The visits of *Centris* species to the flowers of *M. nutans* are aimed at collecting oil, a non-existent reward, which may characterize a form of deceptive pollination. However, upon realizing the absence of oil in *M. nutans* flowers, the bees switch to collecting pollen, a floral resource they also obtain secondarily from Malpighiaceae flowers.

### Is floral mimicry phylogenetically conserved in the genus *Moldenhawera*?

The phenomenon of mimicry is described as a mechanism involving the adaptation of a particular species that potentially increases its reproductive success through the convergence of the floral display with another locally abundant model species. In the case of the genus *Moldenhawera*, the floral morphology is highly conserved, and all species exhibit remarkably similar floral morphologies (Figure 1). As a result, their floral displays collectively bear a striking resemblance to those of Malpighiaceae species. Moreover, the Malpighiaceae family is widely distributed in the Neotropical region and presents a high species diversity in areas where it coexists with *Moldenhawera* species. These combined observations strongly suggest that floral mimicry between *M. nutans* and Malpighiaceae species, as proposed here, may be prevalent across all *Moldenhawera* species, rather than representing an isolated phenomenon specific to a particular species.

If this is indeed the case, it is more parsimonious to hypothesize that the association between *Moldenhawera* species and Malpighiaceae species developed only once during the evolution of the genus and has been maintained in current species due to the advantageous utilization of Centridini bees as pollinators. Centridini bees are a specialized group known for their oil collection behavior, facilitated by oil-collecting structures on the first two pairs of legs (Neff and Simpson 1981; Buchman 1987). This oil, when mixed with pollen, serves as larval food and provides a significantly higher calorie content compared to nectar (Vogel 1983). The dependence of Centridini bees on Malpighiaceae oils for reproduction has been suggested as one of the factors preventing neotropical Malpighiaceae from adapting to other pollinator groups (Anderson 1979, 1990; Vogel 1990; Sigrist 2001). Consequently, in areas where Malpighiaceae species occur, it is likely that oil-collecting Centridini species are also present. Therefore, the pollination of *Moldenhawera* species may result from the phylogenetic conservation of an ancestral pollination system involving floral mimicry with Malpighiaceae species and the exploitation of a highly specialized group of pollinators.

*Moldenhawera* is likely a monophyletic genus, and potential synapomorphies include the presence of T-shaped malpighiaceous trichomes, compound stipules, a heteromorphic androecium with only one fertile stamen on a long filament, and a villous connective (Queiroz et al. 1999; Haston et al. 2003, 2005). Phylogenomic analyses support *Moldenhawera* as the earliest diverging lineage within the Tachigali clade, a group that includes neotropical genera such as *Arapatiella* Rizzini & A. Mattos (2 species), *Diptychandra* Benth. (2), *Jacqueshuberia* Duke (7), and *Tachigali* Aubl. (80–90) (Ringelberg et al. 2022). The genera of this clade exhibit disparate floral morphologies, and no other genera exhibit flowers similar to those of *Moldenhawera*. Therefore, if the floral mimicry of *Moldenhawera* species towards Malpighiaceae is phylogenetically conserved, it is possible to predict that this phenomenon evolved early in the evolutionary history of the genus.

### Other possible candidates: convergence, mimicry, or phylogenetic conservatism?

We found plants exhibiting Malpighiaceae-like floral displays in seven genera of Caesalpinioideae. Among these genera, five are exclusive to the Neotropical region: *Conzattia* Rose (3 species), *Melanoxylum* Schott (1), *Recordoxylon* Ducke (3), *Schizolobium* Vogel (2), and *Moldenhawera* (12). The genus *Bussea* Harms (7 species) is found solely in Africa, while *Peltophorum* (Vogel) Benth. (7) has a pantropical distribution.

The presence of Malpighiaceae-like flowers in Caesalpinioideae species from Africa, Asia, and Australia (such as species of *Bussea* and *Peltophorum*) raises intriguing questions about the underlying processes that have led to the development of this floral display. It is noteworthy that oil-producing Malpighiaceae and oil-collecting Centridini bees are both restricted to the Neotropical region. Consequently, not all instances of similarity in floral display can be attributed to floral mimicry, as they may instead represent examples of convergent evolution or phylogenetic conservatism.

Floral mimicry entails the evolutionary development of floral traits that resemble those of another species, either for deceptive purposes (Batesian mimicry) or for mutual benefits (Müllerian mimicry). In contrast, convergence refers to the independent evolution of similar floral traits in unrelated species due to adaptation to similar ecological conditions. Phylogenetic conservatism, on the other hand, refers to the retention of similar traits within closely related species due to their shared evolutionary history. These concepts offer valuable insights into the diversity and evolution of floral traits within different ecological and evolutionary contexts.

Thus, the examination of various evolutionary hypotheses of floral mimicry requires the fulfillment of distinct criteria. As outlined by Roy and Widmer (1999), for floral similarity between two or more species to be classified as a case of floral mimicry, the following criteria must be satisfied: (1) the species should exhibit significant overlap in their geographical distributions over a substantial period to allow coevolution; (2) they should rely on pollinators for successful seed set; (3) there should be a substantial overlap in their flowering phenology; (4) they must share the same pollinator species, with individual pollinators freely moving between the species; and (5) the observed similarity should confer a significant fitness advantage. Furthermore, we propose an additional criterion, namely, the demonstration that the mimetic species has arisen subsequent to the model species, and in the case of potential floral mimicry with Malpighiaceae, it is important to observe that the visiting behavior of the pollinator bees on the potential mimetic species to be akin to that observed on the model species.

The floral similarity of *Moldenhawera* (discussed above) and *Schizolobium* species appears to fulfill these criteria of floral mimicry. *Schizolobium parahyba* (Vell.) S.F. Blake, in addition to having a floral display very similar to Malpighiaceae species, has one of the stamens adnate and inserted in a furrow of the adaxial petal claw, forming a more robust and resistant structure. Sazima et al. (2009) reported that *Centris labrosa* Friese, 1899 is its main pollinator. This species, along with *Centris varia* (Erichson, 1848), another visitor to the flowers of *S. parahyba*, clasp the reinforced claw of the adaxial petal with their mandibles in a behavior similar to that of these bees when visiting sympatric species of Malpighiaceae. However, these bees do not exhibit oil-collecting behavior; they simply use the claw of the adaxial petal as a support to extend their mouthparts and consume nectar. Sazima et al. (2009) considered this as a case of functional convergence between the claw of the adaxial petal of *S. parahyba* and Malpighiaceae species, although they also considered the possibility of it being a case of floral mimicry. The stem ages of both *Moldenhawera* and *Schizolobium* were estimated at approximately 50 Ma, coinciding with the estimated origin of the *Centris* bees (Figure 3). The lack of accessions from other species within these genera in the reference phylogeny does not allow for the estimation of crown age, but both of them would fall within a period when the oil-producing syndrome of Malpighiaceae and the oil-collecting habit of Centridini bees would be well-established and widespread in the Neotropical region (Martins et al., 2014).

**Figure 3.**
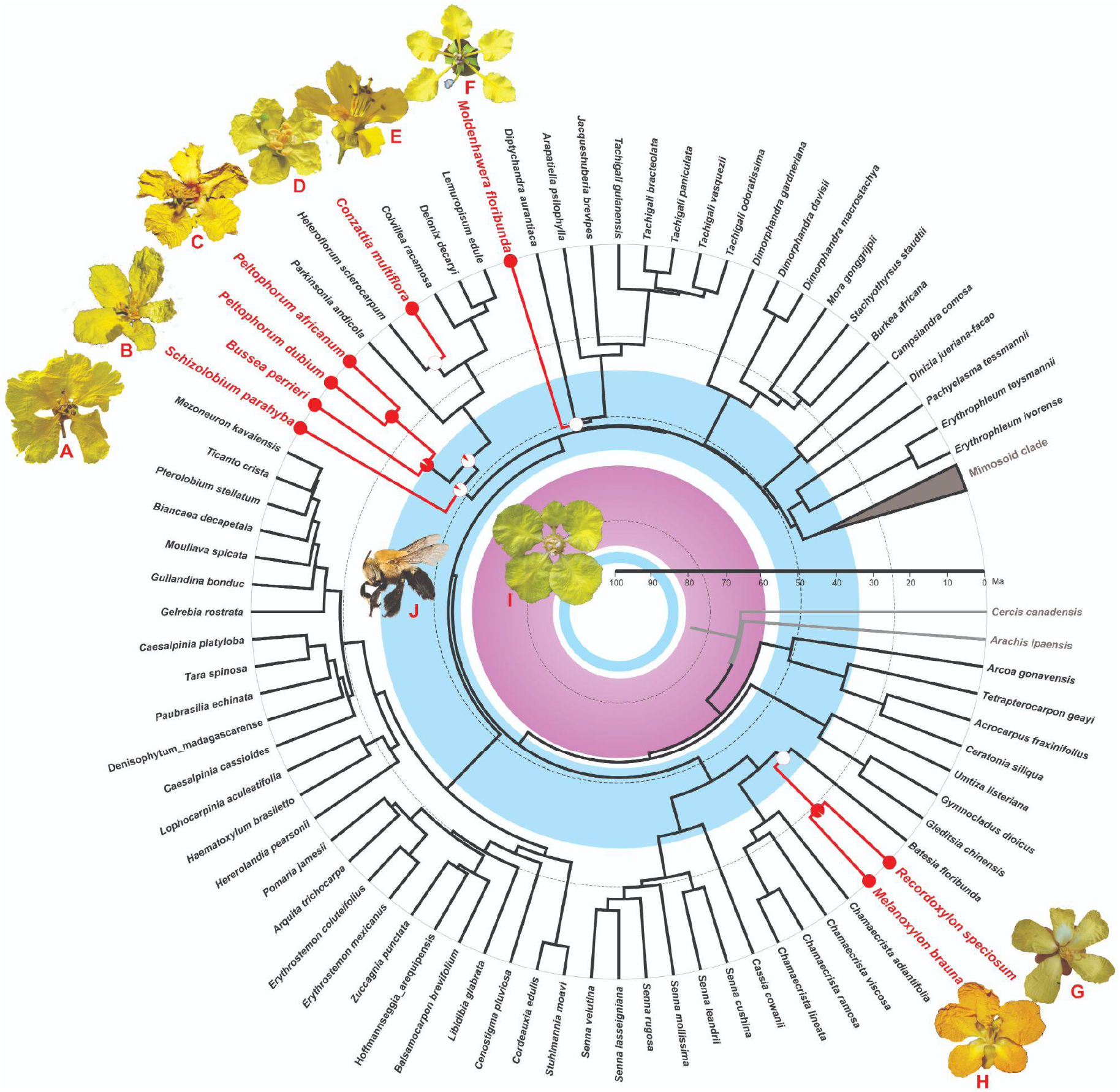
Time-calibrated phylogeny of the subfamily Caesalpinioideae based on a data matrix of 997 nuclear genes presented by Ringelberg et al. (2022b). The highly diverse Mimosoid clade (1,996 terminal taxa) is collapsed. Taxa with Malpighiaceae-like flowers are shown in red. The red dots in some terminals represent taxa with Malpighiaceae-like floral display. The pizzas in some nodes indicate the probability ratio of presence (red) and absence (white) of this type of floral display. The purple circle represents the estimated time of origin of the oil-producing Malpighiaceae, showing the limits of the mean crown age of 59.8 Ma and the mean stem age of 86.1 Ma (Xi et al. 2012). The inner blue circle represents the estimated mean crown age for the appearance of the oil-collecting Centridini bees at about 91 Ma, and the outer blue circle represents the confidence interval for the estimated crown age of the genus *Centris*, ranging from 36 Ma to 59 Ma (mean 46 Ma; Martins et al. 2014). Pictures of the Malpighiaceae-like Caesalpinioideae flowers are from the following species: *Schizolobium parahyba* (Vell.) S.F. Blake (**A**), *Bussea perrieri* R. Vig. (**B**), *Peltophorum dubium* (Spreng.) Taub. (**C**), *Peltophorum africanum* Sond. (**D**); *Conzattia multiflora* (B.L. Rob.) Standl. (**E**); *Moldenhawera blanchetiana* Tul. (**F**); *Recordoxylon speciosum* (Benoist) Gazel ex Barneby (**G**), and *Melanoxylon brauna* Schott (**H**). The flower in the center of the chronogram (**I**) is of the Malpighiaceae species *Peixotoa hispidula* A. Juss. A photo of *Centris nitida* Smith, 1874 is shown in (**J**). Sources of pictures: A: Maurício Mercadante (flickriver.com/photos/mercadanteweb/5081780429/); B: Feno (https://www.inaturalist.org/observations/64635780); C, H: Domingos Cardoso; D: Derek de la Harpe (https://www.inaturalist.org/photos/165849641); E: Colin Hughes; F: Lucas Marinho; G: Glocimar Pereira Silva; I: Rafael F. Almeida; J: The Packer Lab - Bee Tribes of the World (https://en.wikipedia.org/wiki/Centris).

Reproductive fitness tests to demonstrate potential cases of floral mimicry could involve showing that individuals of the mimetic species with floral morphs more similar to the model have higher reproductive success than those with more different morphs (Roy & Widmer, 1999). However, the application of this test is limited to cases where there is significant variation in floral morphology, which was not observed in the studied population of *M. nutans* and does not seem to be the case for the species identified here as having the Malpighiaceae-like floral display. Another type of test would be to demonstrate that populations of the mimetic species have higher reproductive success when occurring sympatrically with the model species compared to in their absence (Roy & Widmer, 1999), a challenging test to perform due to the ubiquity of Malpighiaceae throughout the entire Neotropical region. Due to these challenges, reproductive fitness tests are rarely conducted.

Convergent evolution to similar floral phenotypes does not require overlapping distributions or overlap in flowering phenology, or the demonstration that the similarity is important for reproductive fitness. However, cases of convergence require the demonstration that the similar flowers of unrelated species share the same groups of pollinators and that the floral phenotypes are somehow related to the sensory universe of these pollinators. This corresponds to the classic concept of pollination syndromes (Faegri & van der Pijl 1980) and should serve as the first step in determining the other evolutionary processes discussed here. In turn, cases of phylogenetic inertia represent the maintenance of a floral phenotype resulting from floral mimicry throughout the lineage, even in the absence of the model species.

Differentiating cases of phylogenetic conservatism from simple morphological convergence in potential Malpighiaceae-like floral displays can be achieved by reconstructing this character state in the ancestor of a group in which some species with this phenotype no longer interact with these bees. This may be the case for Old World species of the genera *Bussea* and *Peltophorum*, which belong to lineages whose ancestors have been reconstructed as having Malpighiaceae-like floral displays. Unfortunately, we did not find empirical studies on the pollination of species of these genera, and the sampling of species from these genera in the phylogeny (three out of 14 species) does not allow for more elaborate considerations.

### Concluding remarks

Our hypothesis posits that *Moldenhawera nutans* exhibits floral mimicry towards Malpighiaceae flowers, substantiated by several lines of evidence. First, we observed a remarkable resemblance in the floral display between *M. nutans* and Malpighiaceae species. Second, both *M. nutans* and Malpighiaceae flowers were visited by the same species of oil-collecting *Centris* bees, implying a shared pollinator preference. Furthermore, despite the absence of oil production in *M. nutans*, we documented the peculiar behavior of *Centris* bees initially attempting oil collection, subsequently adapting to pollen collection upon realizing the absence of oil.

This study opens an intriguing avenue for exploring other potential cases of floral mimicry between species of Caesalpinioideae and Malpighiaceae. Other potential cases of floral mimicry with Malpighiaceae species have been analyzed, but with the current data, it is not possible to hypothesize whether they truly represent floral mimicry or are the result of convergence or phylogenetic inertia. If they are indeed cases of true floral mimicry, we predict that the oil-collecting pollinator species should exhibit similar visiting behavior to that observed in Malpighiaceae flowers, even in the absence of oil in Caesalpinioideae flowers.

It would be interesting to integrate phylogenetic information with studies on pollination in the mentioned genera of Caesalpinioideae. If floral mimicry with Malpighiaceae is an ancestral and conserved mechanism throughout the phylogeny of a genus, we would expect to find the same types of pollinators with the same stereotyped behavior in the current species of the genus, along with a strong similarity in floral displays, as suggested by the striking resemblance of *Moldenhawera* flowers.

## Acknowledgments

We would like to thank Brigitte Marazzi, Domingos Cardoso, and Carolina L. Ribeiro for their valuable comments on an earlier version of the manuscript, and Jens Ringelberg for kindly sharing the chronogram of the Caesalpinioideae subfamily. The photos illustrating this article were kindly provided by Domingos Cardoso, Lucas Marinho, Alex Popovkin, Glocimar P. Silva, Rubens Queiroz, Colin E. Hughes, Rafael F. Almeida, Marco Pellegrini, and Cláudio N. Fraga. LPQ studies on Leguminosae are supported by a PQ-1A grant from CNPq (Process No. 305230/2021-2).

## References

Almeida RF (2020a) Peixotoa in Flora e Funga do Brasil. Jardim Botânico do Rio de Janeiro. Available at <https://floradobrasil.jbrj.gov.br/FB8933>. Acessed 20 May 2023

Almeida RF (2020b) Stigmaphyllon in Flora e Funga do Brasil. Jardim Botânico do Rio de Janeiro. Available at <https://floradobrasil.jbrj.gov.br/FB8939>. Acessed 20 May 2023

Anderson WR (1979) Floral conservatism in Neotropical Malpighiaceae. Biotropica 11: 219–223. DOI 10.2307/2388042.

Anderson WR (1990) The origin of Malpighiaceae - the evidence from morphology. Memoirs of the New York Botanical Garden 64: 219–224.

Arroyo MTK (1981) Breeding systems and pollination biology in Leguminosae. Pp 723–769 in Polhill RM, Raven PH, Eds., Advances in Legume Systematics, Part 2, Royal Botanic Garden, Kew.

Brown JH, Kodric-Brown A (1979) Convergence, competition and mimicry in a temperate community of hummingbird-pollinated flowers. Ecology 60: 1022–1035. DOI 10.2307/1936870.

Buchmann LS (1983) Buzz pollination in Angiosperms. Pp 73–113 in Jones CE, Little RJ, Eds., Handbook of experimental pollination biology. Scientific and Academic Editions, New York.

Buchmann SL (1987) The ecology of oil flowers and their bees. Annual Review of Ecology and Systematics 18: 343–369. DOI http://www.jstor.org/stable/2097136.

Costa CBN, Costa JAS, Ramalho M (2006) Biologia reprodutiva de espécies simpátricas de Malpighiaceae em dunas costeiras da Bahia. Brasil. Revista Brasileira de Botânica 29: 103–114. DOI: 10.1590/S0100-84042006000100010.

Dafni A (1984) Mimicry and deception in pollination. Annual Review of Ecology and Systematics 15: 259–278. DOI: http://www.jstor.org/stable/2096949.

Davis CC, Anderson WR, Donoghue MJ (2001) Phylogeny of Malpighiaceae: Evidence from Chloroplast ndhF and trnL-F Nucleotide Sequences. American Journal of Botany 88: 1830–1846. DOI 10.2307/3558360.

Davis CC, Fritsch PW, Bell CD, Mathews S (2004) High latitude Tertiary migrations of an exclusively tropical clade: evidence from Malpighiaceae. International Journal of Plant Science 165: S107–S121. DOI: 10.1086/383337.

Faegri K, van der Pijl L. (1980) [1966]. Principles of Pollination Ecology (3rd ed.). Elsevier.

Gagnon E, Bruneau A, Hughes CE, Queiroz LP, Lewis GP (2016) A new generic system for the pantropical Caesalpinia group (Leguminosae). PhytoKeys 71: 1–160. DOI: 10.3897/phytokeys.71.9203.

Haston EM, Lewis GP, Hawkins JA (2003) A phylogenetic investigation of the Peltophorum group (Caesalpinieae: Leguminosae). Pp 149–159 in: Klitgaard BB, Bruneau A., Eds, Advances in Legume Systematics, part 10. Royal Botanic Gardens, Kew.

Haston EM, Lewis GP, Hawkins JA (2005) A phylogenetic reappraisal of the Peltophorum group (Caesalpinieae: Leguminosae) based on the chloroplast trnL-F, rbcL and rps16 sequence data. American Journal of Botany 92: 1359–1371. DOI: 10.3732/ajb.92.8.1359

Johnson SD (1994) Evidence for Batesian mimicry in a butterfly-pollinated orchid. Biological journal of the Linnean Society 53: 91–104. DOI: 10.1111/j.1095-8312.1994.tb01003.x

Johnson SD, Alexandersson R, Linder H. (2003) Experimental and phylogenetic evidence for floral mimicry in a guild of fly-pollinated plants. Biological Journal of the Linnean Society 80: 289–304. DOI: 10.1046/j.1095-8312.2003.00236.x

Little RJ (1983) A review of floral food deception mimicries with comments on floral mutualism. Pp. 294–309 in Jones CE, Little RJ, Eds., Handbook of Experimental Pollination Biology. Van Nostrand Reinhold.

Macior, L.W. (1971) Co-evolution of plants and animals – systematic insights from plant–insect interactions. Taxon 20: 17–28. DOI: 10.2307/1218530

Maddison WP, Maddison DR (2023) Mesquite: a modular system for evolutionary analysis. Version 3.81 http://www.mesquiteproject.org.

Martins AC, Melo GAR, Renner SS (2014) The corbiculate bees arose from New World oil-collecting bees: Implications for the origin of pollen baskets. Molecular Phylogenetics and Evolution 80: 88–94. DOI: 10.1016/j.ympev.2014.07.003

Neff JL, Simpson B. (1981) Oil-collecting structures in Anthophoridae (Hymenoptera): morphology, function and use in systematics. Journal of the Kansas Entomological Society 54: 95–123. DOI: http://www.jstor.org/stable/25084137

Papadopulos AST, Powell MP, Pupulin F, Warner J, Hawkins JA, Salamin N, Chittka L, Williams NH, Whitten WM, Loader D, Valente LM, Chase MW, Savolainen V (2013) Convergent evolution of floral signals underlies the success of Neotropical orchids. Proc. R. Soc. B 280: 20130960. DOI: 10.1098/rspb.2013.0960

Peter CI, Johnson SD (2008) Mimics and magnets: the importance of color and ecological facilitation in floral deception. Ecology 89: 1583–1595. DOI: http://www.jstor.org/stable/27650665

Proctor M, Yeo P (1972) The Pollination of Flowers, Taplinger, New York, USA.

Queiroz LP, Lewis GP, Allkin R (1999) A revision of the genus Moldenhawera Schrad. (Leguminosae–Caesalpinioideae). Kew Bulletin 54: 817–852. DOI: 10.2307/4111163

Ringelberg JJ, Koenen EJM, Iganci JR, Queiroz LP, Murphy DJ, Gaudeul M, Bruneau A, Luckow M, Lewis GP, Hughes CE (2022) Phylogenomic analysis of 997 nuclear genes reveals the need for extensive generic re-delimitation in Caesalpinioideae (Leguminosae). In: Hughes CE, Queiroz LP, Lewis GP, Eds, Advances in Legume Systematics 14. Classification of Caesalpinioideae Part 1: New generic delimitations. PhytoKeys 205: 3–58. DOI: 10.3897/phytokeys.205.85866

Ringelberg JJ, Koenen EJM, Sauter B, Aebli A, Rando JG, Iganci JR, Queiroz LP, Murphy BJ, Gaudeul M, Bruneau A, Luckow M, Lewis GP, Miller JT, Simon MF, Jordão LSB, Morales M, Bailey CD, Nageswara-Rao M, Nicholls JA, Loiseau O, Pennington RT, Dexter KG, Zimmermann NR, Hughes CE (2023) Precipitation is the main axis of tropical plant phylogenetic turnover across space and time. Science Advances 9 (7): eade4954. DOI: 10.1126/sciadv.ade4954

Roy BA, Widmer A (1999) Floral mimicry: a fascinating yet poorly understood phenomenon. Trends in Plant Science 4: 325–330. DOI: 10.1016/s1360-1385(99)01445-4

Sazima M, Sazima I (1989) Oil-gathering bees visit flowers of eglandular morphs of the oil-producing Malpighiaceae. Botanica Acta 102: 106–111. DOI: 10.1111/j.1438-8677.1989.tb00073.x

Sazima I, Pinheiro M, Sazima M (2009) A presumed case of functional convergence between the flowers of Schizolobium parahyba (Fabaceae) and species of Malpighiaceae. Plant Systematics and Evolution 281: 247–250. DOI 10.1007/s00606-009-0193-5

Schemske DW (1981) Floral convergence and pollinator sharing in two bee-pollinated tropical herbs. Ecology 62: 946–954. DOI: 10.2307/1936993

Sigrist MR (2001) Reproductive biology of twelve sympatric species of Malpighiaceae in semideciduous forest in southeastern Brazil. Universidade Estadual de Campinas, PhD thesis.

Sigrist MR, Sazima M (2004) Pollination and reproductive biology of twelve species of neotropical Malpighiaceae: stigma morphology and its implications for the breeding system. Annals of Botany 94: 1–9. DOI: 10.1093/aob/mch108

Thomson JD (1981) Spatial and temporal components of resource assessment by flower-feeding insects. Journal of Animal Ecology 50, 49–59. DOI: 10.2307/4030

Viana BF, Kleinert AMP (2006) Structure of bee-flower system in the coastal sand dune of Abaeté, Northeast of Brazil. Revista Brasileira de Entomologia 50: 53–63. DOI: 10.1590/S0085-56262006000100008

Viana BF, Silva FO, Kleinert AMP (2006) A flora apícola de uma área restrita de dunas litorâneas, Abaeté, Salvador, Bahia. Revista Brasileira de Botânica 29: 13–25. DOI: 10.1590/S0100-84042006000100003

Vivas CV, Queiroz LP (2020) Moldenhawera in Flora e Funga do Brasil. Jardim Botânico do Rio de Janeiro. Available at <https://floradobrasil.jbrj.gov.br/FB28148>. Accessed 20 May 2023.

Vivas CV, Gaiotto FA, Queiroz LP (2015) A new species of Moldenhawera (Leguminosae) from Brazilian Atlantic Forest. Phytotaxa 233: 85–89. DOI: 10.11646/phytotaxa.233.1.7

Vivas CV, Souza G, Gaiotto FA, Queiroz LP (2019) Moldenhawera congestiflora: a new species of Leguminosae from the Brazilian Atlantic Forest. Phytotaxa 399: 285–290. DOI: 10.11646/phytotaxa.399.4.4

Vogel S (1974) Ölblumen und ölsammelnde Bienen. Tropische und subtropische Pflanzenwelt 7: 283–547.

Vogel S (1983) Ecophysiology of zoophilic pollination. Pp. 559–624 in Lange OL, Nobel PS, Osmond CB, Zeigler H, Eds., Encyclopedia of plant physiology, new series 12C: Physiological Plant Ecology III. Berlin, Springer.

Vogel S (1990) History of the Malpighiaceae in the light of pollination ecology. Memoirs of the New York Botanical Garden 55: 341–362.

Westerkamp C (2004) Ricochet pollination in cassias - and how bees explain enantiostyly. Preliminary communication. Pp. 225–230 in Freitas BM, Pereira JOP, Org., Solitary Bees. Conservation, Rearing and Management for Pollination. Fortaleza: Imprensa Universitária.

Xi Z, Ruhfel BR, Schaefer H, Amorim AM, Sugumaran M, Wurdack KJ, Endress PK, Matthews M, Stevens PF, Mathews S, Davis CC (2012) Phylogenomics and a posteriori data partitioning resolve the Cretaceous angiosperm radiation in Malpighiales. Proceedings of the National Academy of Sciences 109: 17519–17524. DOI: 10.1073/pnas.1205818109

